# Real-time halo correction in phase contrast imaging

**DOI:** 10.1101/227025

**Authors:** Mikhail E. Kandel, Michael Fanous, Catherine Best-Popescu, Gabriel Popescu

## Abstract

As a label-free, nondestructive method, phase contrast is by far the most popular microscopy technique for routine inspection of cell cultures. Yet, features of interest such as extensions near cell bodies are often obscured by a glow, which came to be known as the halo. Advances in modeling image formation have shown that this artifact is due to the limited spatial coherence of the illumination. Yet, the same incoherent illumination is responsible for superior sensitivity to fine details in the phase contrast geometry. Thus, there exists a trade-off between high-detail (incoherent) and low-detail (coherent) imaging systems. In this work, we propose a method to break this dichotomy, by carefully mixing corrected low-frequency and high-frequency data in a way that eliminates the edge effect. Specifically, our technique is able to remove halo artifacts at video rates, requiring no manual interaction or *a priori* point spread function measurements. To validate our approach, we imaged standard spherical beads, sperm cells, tissue slices, and red blood cells. We demonstrate the real-time operation with a time evolution study of adherent neuron cultures whose neurites are revealed by our halo correction. We show that with our novel technique, we can quantify cell growth in large populations, without the need for thresholds and calibration.

## 1. Introduction

In developing phase contrast microscopy (PCM)[1], Zernike employed Abbe’s revolutionary concept in imaging theory: the image is a result of interference between the incident and the scattered light, which is to say, the image is an interferogram[2]. Thus, the problem of enhancing the contrast in an image of a transparent specimen becomes equivalent to increasing the contrast in an interferogram. By developing a spatial filter that both attenuates and phase-shifts the incident light by ***π***/2 with respect to the scattered light, Zernike obtained drastically improved contrast even from complete phase objects, such as those specimens exhibiting negligible absorption. This achievement triggered the broad adoption of PCM by most cell biology labs, as the instrument offers fast, nondestructive visual inspection of unlabeled cells.

Despite its massive success, PCM is not without limitations. Since the intensity image depends on four variables: the intensity of the incident light, the intensity of the scattered light, the sine, and cosine of their phase difference, PCM’s use is limited to *qualitative* inspection rather than *quantitative* assays. Furthermore, because the incident light is not a perfect plane wave, it contains non-zero frequency components that get inadvertently phase shifted by ***π***/2, and thus, interfere with the scattered light. These components result in contours of high-contrast intensity changes, especially around the edges of the object, commonly referred to as “halo”. Previous work has been dedicated to eliminating the halo problem. Double-path systems were once popular[3, 4], although such systems are known to suffer from poor temporal phase stability, constraining time-lapse observation. A more popular class of solutions involves changing the illumination to a spatially coherent field[5-8], which involves modifying the instrument and, thus, reducing its compatibility with existing facilities. Similarly, it is possible to start from a commercial phase contrast microscope and simply reduce the area of the illumination ring, thus improving the spatial coherence of the illumination leading to a reduction in halo[9, 10]. Perhaps the most immediate way to increase the coherence of the imaging field is to switch to a laser line illumination[11-13]. Yet, when the coherence area becomes much larger than the field of view, there is a notable reduction in image quality due to laser speckle and other coherence-induced artifacts, such as spurious fringes, diffraction rings, etc[12].

The second class of approaches involves pure numerical processing, which are particularly popular in phenotypic screening applications[14]. For example, in references [15, 16] the halo was used to improve the algorithm’s ability to discriminate between cells, yet in[17] it was noted that halos obscure morphological features. Invariant to the author’s cell segmentation strategy, the resulting cell perimeter coordinates must be further refined in post-processing[18, 19]. Finally, a method that combines both hardware and numerical processing was proposed in Ref. [20]. Using a full description of image formation with partially coherent light, this approach relies on iterative deconvolution steps to invert a nonlinear image formation model. While successful for certain categories of sample, the approach suffers from poor numeric convergence leading to long computation times, impractical for real-time operation and high-throughput assays.

In this paper, we describe a new, real-time, approach for removing halos in existing phase contrast microscopy. Our method uses an optical module that attaches to the output of an otherwise unmodified microscope (Fig. 1(a)). This module allows us to collect four interference images, which are used to decouple the unknowns in a per-pixel interferometric measurement and obtain a map of optical path length shifts[21]. As this quantitative phase image is a faithful measurement of the image field, it, too, contains the halo artifact (Fig. 1(b)). Using numerical processing that involves a spatial Hilbert transform; we show that the edge artifacts are effectively removed. The main advantages of this approach are: 1) the halo removal procedure runs in real-time 2) the method combined with the SLIM hardware can be used to augment existing phase contrast microscopes, and 3) the fidelity of the quantitative phase map improves, enabling quantitative measurements on cells and tissues. We show that this procedure can be applied to other common-path imaging methods that suffer from artifacts due to partially coherent illumination. We demonstrate our technique on standard samples, a variety of live cells, as well as time-lapse measurements of neuronal growth that would be otherwise difficult to measure without our method.

**Fig. 1.**
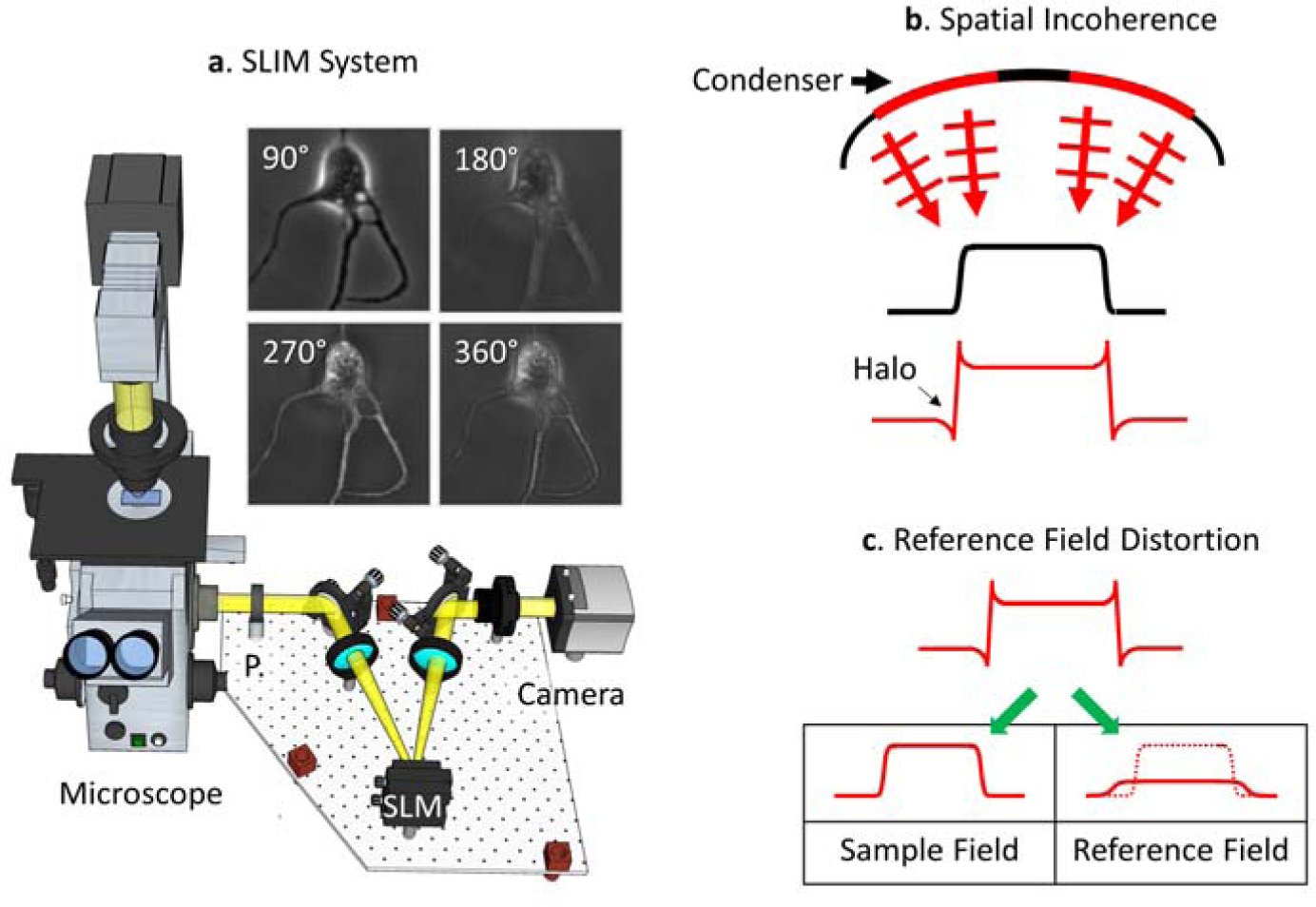
A halo forms when the reference field is distorted by a low-pass version of the sample field. (a) Measurement of the halo-artifact was performed using a four-frame shifting interferometer (SLIM) attached to a phase-contrast microscope. From four orthogonal interferograms (90°, 180°, 270°, 360°) it is possible to uniquely determine the phase at each pixel. (b) Due to its wide illumination aperture (h_o_, red), the phase contrast condenser introduces a spread in spatial frequencies,leading to unwanted glows in the measured field (arg [Γ]) when compared to the object, the condenser introduces a spread in arg [T]). (c) Due to the spread in illuminating spatial frequencies, the reference field is not flat, but rather contains a distorted, low-frequency replicate of the sample field.

## 2. Materials and methods

### Microscopy

Our method can be implemented with any phase-sensitive imaging technique. However, we choose to use quantitative phase imaging[22] (QPI), which enables objective measurements of the phase shift in a reproducible, system-independent manner. Recently, such systems have found fertile ground in studies of cellular morphology[23-31], plate reader style screening of whole cell populations[24, 32-35], and histopathology sections[36, 37].

Phi Optics, Inc., Fig. 1(a)) which is highly sensitive to phase shifts from fine object details[38]. The module was coupled to a commercial PCM (Axio Observer Z1, Zeiss). The components used in our add-on module are a small contribution to the total cost of ownership of a fully automated commercial microscope. The system essentially performs four-frame phase shifting interferometry[39] on the phase-contrast microscope field. After numerical processing, the four phase-contrast frames yield a measurement of the phase shift along the optical path at each pixel. The precise operation of the SLIM module is described elsewhere[38]. To demonstrate the broad applicability of our halo removal technique, the RBC in Fig. 5(d) was measured on an off-axis common-path white light interferometer[40], which also suffers from halo artifacts[41].

### Sample preparation

We imaged a variety of samples, as illustrated in Fig. 5. The HeLa cell culture (Fig. 4 (b)) was prepared following reference [42]. Polystyrene beads (Fig. 4(b) & Fig. 5(a)) are a common metrology sample used to verify the accuracy of phase sensitive microscopes. The beads were suspended between two coverslips and imaged with a 40x/0.75 objective in oil immersion media with a well-known refractive index (Zeiss, n=1.518). The oil immersion is used to avoid “phase wrapping”[43]. The spermatozoon (Fig. 5(b)) was obtained from freshly thawed bull semen, which after washing, was imaged in a phosphate-buffered saline solution. The tissue microarray (Fig. 5(c)) was prepared according to the protocol in reference[44]. Following reference[45], the RBC (Fig. 5(d)) was imaged in a PBS solution spaced between two coverslips.

**Fig. 4.**
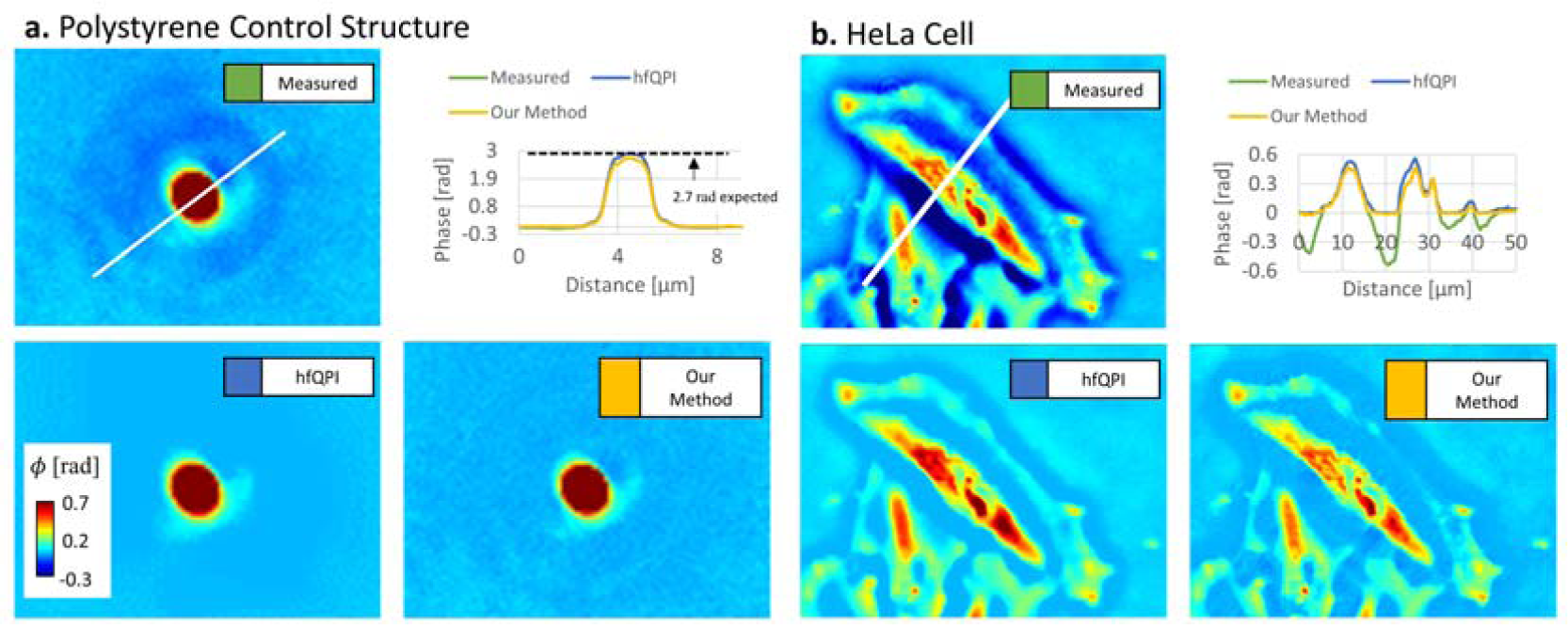
To compare to previous, not real-time, halo removal efforts - “hfQPI” in Nguyen et al. (2017) -we imaged a polystyrene bead (a) as well as adherent cells (b). (a) When Halo removal applied to a 3-μm polystyrene bead shows a line profile fitting well with the expected peak phase value of 2.7 radians. (b) Applying our method to a more complicated structure such as a cluster of HeLa cells, we note that for this category of sample, results are qualitatively similar and stress that our approach is real-time. Data acquired with 20x/0.3.

**Fig. 5:**
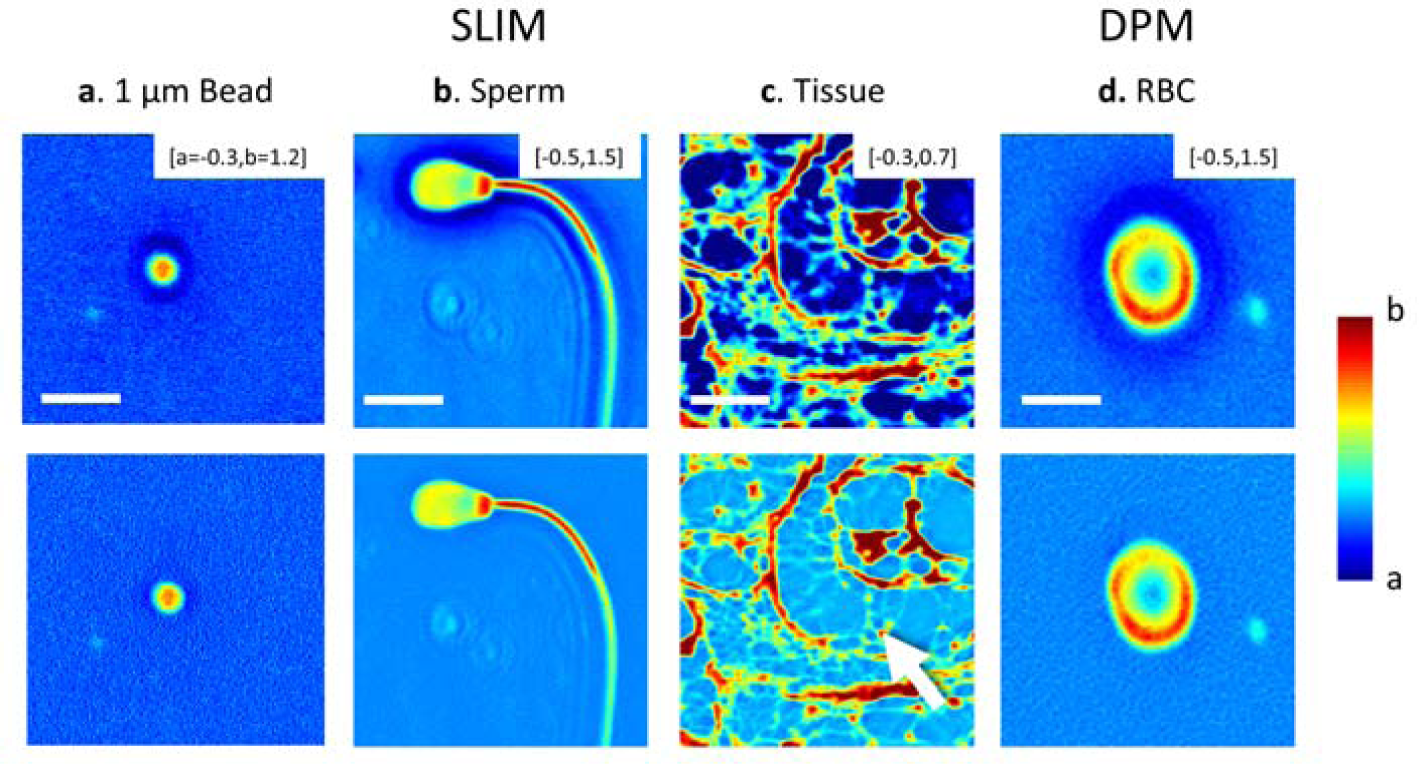
Halo removal reveals submerged high-frequency content. Because our halo removal routine includes the existing phase map, the procedure preserves the shape of control structures such as the 1 μm polystyrene bead (white bar = 5 μm). We note that there is little change in the image as the structure does not exhibit a significant halo. The prominent halo around the sperm cell is removed after application of our technique (white bar = 12 μm). Notably, in continuous samples, such as surgically resected tissue (white bar = 5 μm), the halo often obscures significant details such as the fibers separating individual cells in gland structures. While images in this paper were acquired with a SLIM style add-on module, the procedure is also applicable to other common-path systems such as the DPM microscope (RBC, white bar = 6 μm). Phase maps are displayed in a range from a, b.In this work, we use a phase contrast based phase-shifting interferometer (CellVista SLIM Pro module,

The postnatal mice neurons (P0-P1, C57BL/6) were prepared as described in[26]. After rapidly thawing, the neurons were plated on poly-d-lysine-coated 35 mm glass Petri dishes at a low-density (∼65 cells per mm^2^) and grown at 37°C, in the presence of 5% CO_2_, in standard maintenance media Neurobasal^®^ 1% 200 mM glutamine and 1% penicillin/streptomycin. Reagents were obtained from Invitrogen. Time-lapse observation was performed after 5 days *in vitro*. Half the media was aspirated and replaced with fresh maintenance media warmed to 37°C immediately before imaging.

## 3. Results and discussion

We use a Hilbert transform approach to correct the high-frequency data, effectively generating halo-free images. In essence, we filter the halo with directional derivatives, performing a 1D integration for each integration, and merge the resulting images, in the spatial domain, with undistorted content. This procedure effectively preserves the “correct” parts of the phase map and fixes the incorrect ones. The method is motivated by the observations in Fig. 2. Thus, the halo appears around the edges of the cell as a thick, negative phase value shadow (Fig. 2(a), red arrows). Yet, the halo is not a significant issue for isolated, small, structures like the neurite extension in Fig. 2(a) (green arrow). Thus, the image’s spatial frequency content can be broadly divided into “preserved high-frequencies” and “distorted low-frequencies” (Fig. 2(b)).

**Fig. 2.**
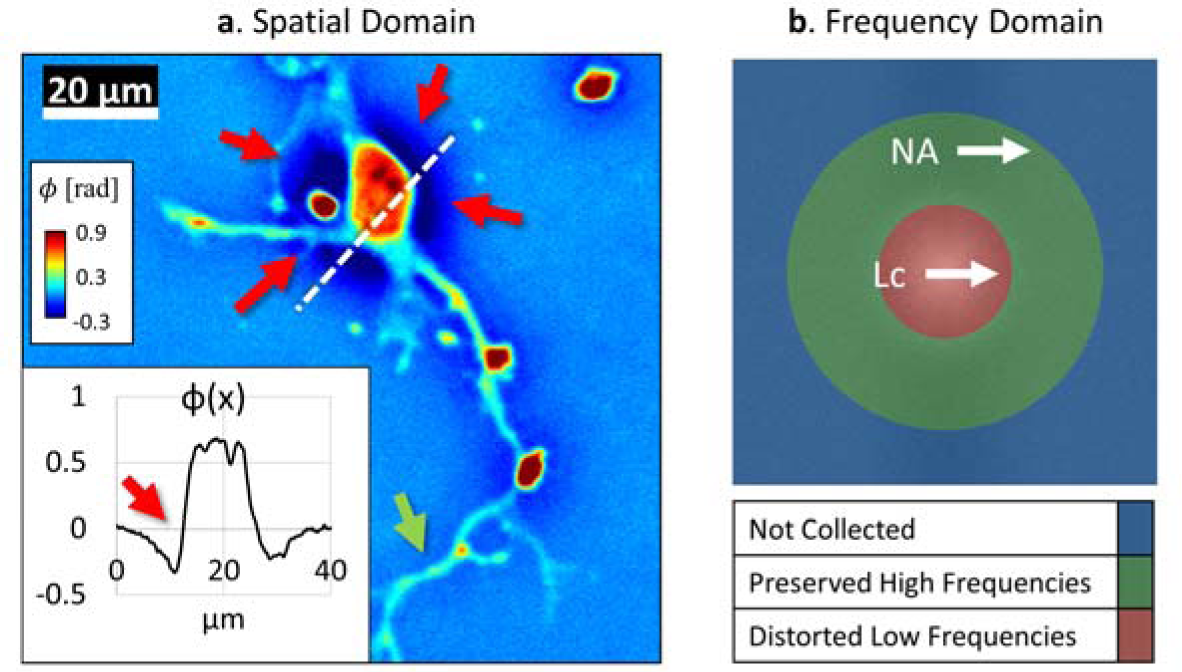
The halo appears as a shadow that disproportionately affects low-frequency content. (a) A typical phase map (inset profile) the halo appears as a negative glow (phase delay) around the cell body (red arrows). The defect is negligible for isolated, small, structures typical of neuronal extensions (green arrow). (b) In the frequency domain, the image divides into “preserved high-frequencies” (green), and “distorted low-frequencies”.

The algorithm proceeds by taking a directional derivative of the frequency bands corresponding to objects smaller than the coherence radius. When applied to a phase map, this procedure yields an image similar to simulated differential interference contrast (DIC). This “relief” style image contains significantly less halo compared to the input image, as it is essentially a high-pass filtered gradient. To remove the gradient and highlight the edges otherwise submerged by the halo, we use a Hilbert transform to perform integration along the direction of the derivative. In the first approximation, the measured phase shift, *φ_m_*, can be described as the ideal phase, *φ*, from which a low-frequency version, 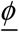, is subtracted[20],

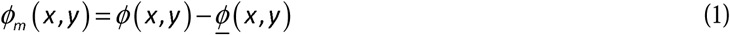

Equation 1 describes how portions of the measured image exhibit negative values. The key idea of our approach is to realize that since 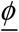 is a smooth function and, thus, its spatial derivative is negligible. Taking the derivative along a direction, say ***X***, we obtain a measured quantity that is not significantly affected by the halo,

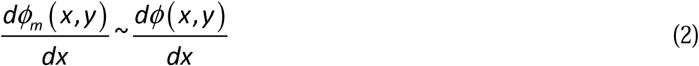

In order to integrate and obtain back the quantitative phase, we use the Hilbert transform along ***X***. The Hilbert transform is a common tool in off-axis holography [46-48] and has been used in the context of differential interference contrast microscopy (DIC) as well [49, 50].

Taking the Hilbert transform of Eq. 2 yields

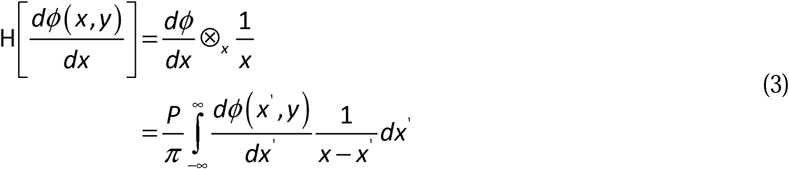

In Eq. 3, ***P*** denotes a principal value integral and ⊗*_x_* stands for the convolution operation along ***X***. Integrating Eq. 3 by parts, we obtain

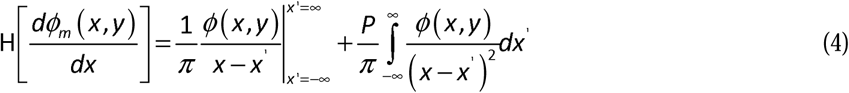

Note that the first term on the right-hand side of Eq. 4 vanishes and the second term amounts to a convolution operation, thus

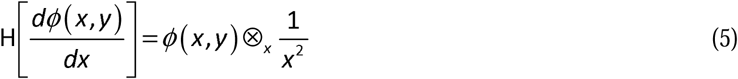

Equation 5 indicates that applying a Hilbert transform to the phase derivative yields the phase image itself, except, it is convolved with a kernel 1 / ***X***^2^. This means that the phase reconstruction is very accurate at short scales when the kernel approaches a delta-function. Therefore, to make sure that large-scale features are not washed out by this convolution, we compare each pixel in the reconstructed phase with the original measurement and keep the maximum. With this procedure, we ensure that the Hilbert transform can only improve the image. Since the Hilbert, transform along one direction is most effective at removing the halo at the edge perpendicular to the direction of the derivative; we compute the Hilbert transform along multiple directions and choose at each point the maximum phase value. This way, in essence, we identify the dominant edge direction at each point in the image and remove the respective halo artifact.

For maximum computation speed, we perform both the spatial derivative, and Hilbert transforms simultaneously (Fig. 3), in the frequency domain, as

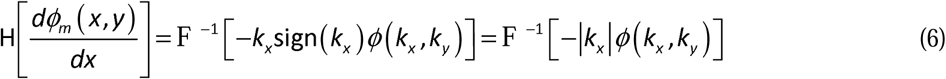

To account for different objectives and illumination sources, spatial frequencies corresponding to those above the coherence length are completely preserved (Fig. 3, 1/***L***_*c*_ shoulder in the filter profile). We emphasize that this filter choice is independent of the sample and only needs to be made once for each objective. We choose ***L***_*c*_= **2 *μm*** and stress that no qualitative differences are observed within an order of magnitude. Thus, Eq. 6 becomes,

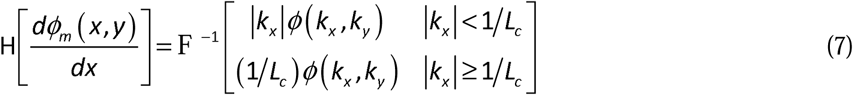

**Fig. 3.**
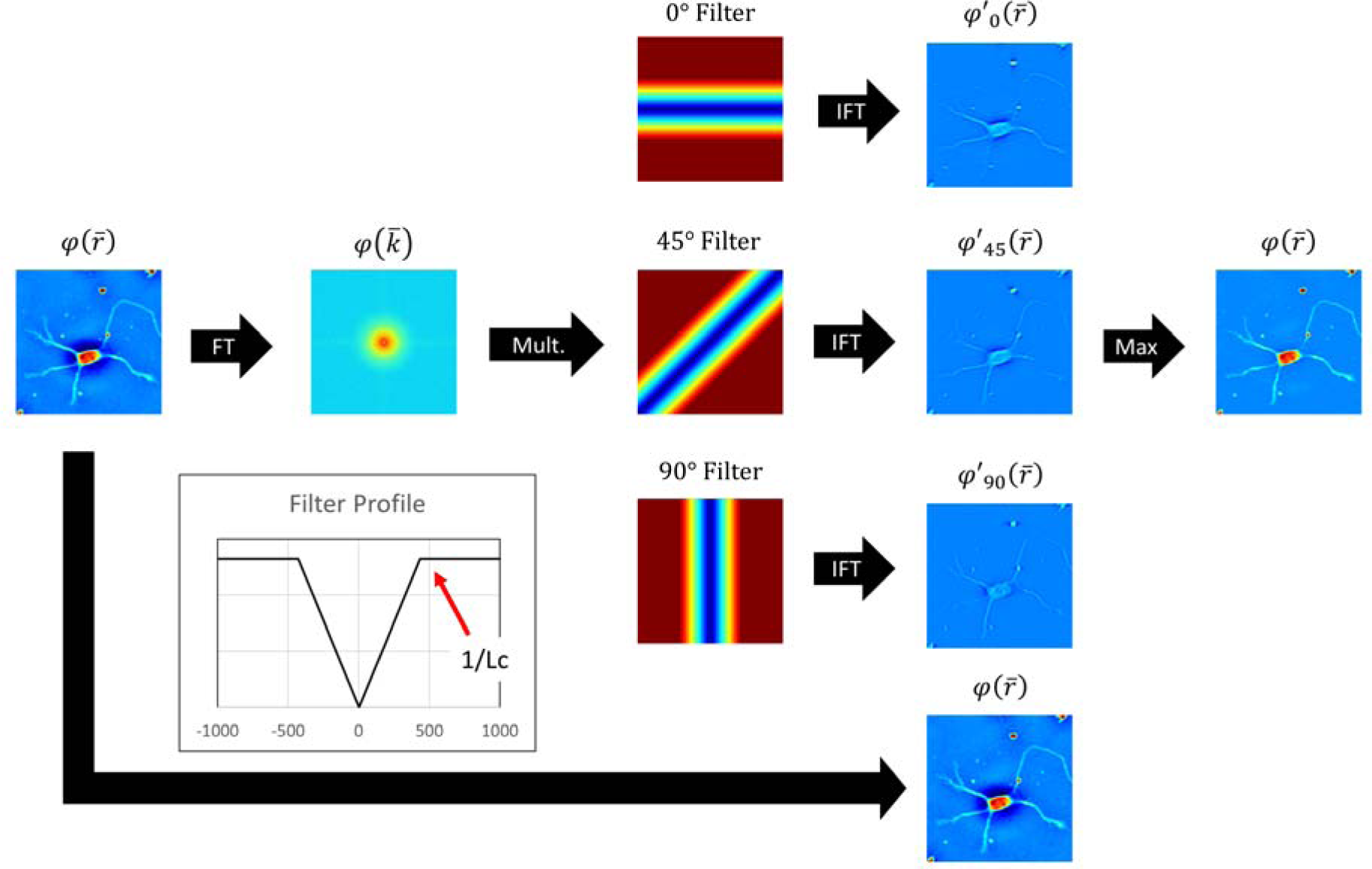
A direct (non-iterative) algorithm to remove halo artifacts using the Hilbert transform. Directional filters are applied to the frequency domain representation of the image. The frequency content corresponding to fine details unaffected by the halo ( those greater than 1/L_c) are allowed to pass un-perturbed. For the low frequencies content affected by the halo we apply a filter corresponding to a derivative combined with a signum function. In our implementation we use three such directions, and take the imaginary part of the inverse Fourier transform. These three directional images, as well as the original image are then merged by taking a pixel wise maximum of the values, such that in areas without a halo there is no change in pixel value.

We found that three operations along different directions were sufficient for a good reconstruction (Fig. 3, along 0°, 45°, 90°). Due to the high-speed of our GPU-based implementation, typical computation times are under 40 ms and dominated by the Fourier transform step (GeForce GTX 970M, 4 MP phase map). Visualization 1-4 (movies) show representative videos of the system’s operation. Visualization 4 (movie) illustrates the real-time operation and that the computation time is less than the typical time to acquire a frame. In practice, this means the proposed algorithm is completely masked by the acquisition process [44]. Fig. 4 shows a comparison between the proposed algorithm and a previous efforts [20], a runtime comparison is omitted as our method is implemented as an optimized GPU code, while the latter is written in MATLAB and runs on the CPU.

Fig. 5 shows that our demodulation routine is applicable to a wide category of samples. First, we tested our method on standard samples. The 1 μm bead in Fig. 5 is a common metrology sample used to verify the accuracy of phase sensitive microscopes. The polystyrene beads were suspended between two coverslips and imaged with a 40x/0.75 objective in oil immersion. Importantly, the round shaped structure remains completely unchanged after restoration.

The bovine spermatozoon shown in Fig. 5 has a dense halo artifact around the edges of the head region (40x/0.75). After processing, the halo disappears and the negative values are removed. Unlike the sperm cell in Fig. 5(b), the 4 μm thick slice of histopathological prostate tissue in Fig. 5(c) forms a continuous sheet spanning a large area of the microscope slide. The halo surrounds the gland wall (white arrow). This artifact effectively submerges fine extracellular structures under a large negative valued shadow. After applying our method, we can clearly see the cell walls of the small cells that make up the many lobbed acinary shaped gland (Fig. 5, white arrows). Importantly, we can correct the halo while preserving the relative thickness of gland wall. The ability to preserve low-frequency data while recovering high-frequency information is particularly important for studying adherent cells with complicated morphology.

To demonstrate the wide-ranging applications of our method, we performed halo removal on a red blood cell acquired with white light diffraction phase microscopy (wDPM[40], 40x/0.75NA objective). As wDPM is a self-referenced interferometer operating under broadband illumination, it, too, can display halo artifacts. As shown in Fig. 5, the halo from this very different microscope is also removed.

To validate our technique as a method to study cell growth and proliferation, we performed time-lapse imaging of a whole 35 mm petri dish over the course of 29 hours (Fig. 6, 20x/0.3NA objective). This is accomplished by using the automated imaging technique described in[44], where the image consists of a large number of mosaic tiles that are assembled and aligned to form a two gigapixel time-lapse sequence. Once assembled the sequence can be inspected with a “deep zoom” style image viewer (such as TrakEM2[51]). By manually counting representative fields of view, we estimate that the dish contained 64 thousand neurons and 45 thousand well-pronounced neurites.

**Fig. 6.**
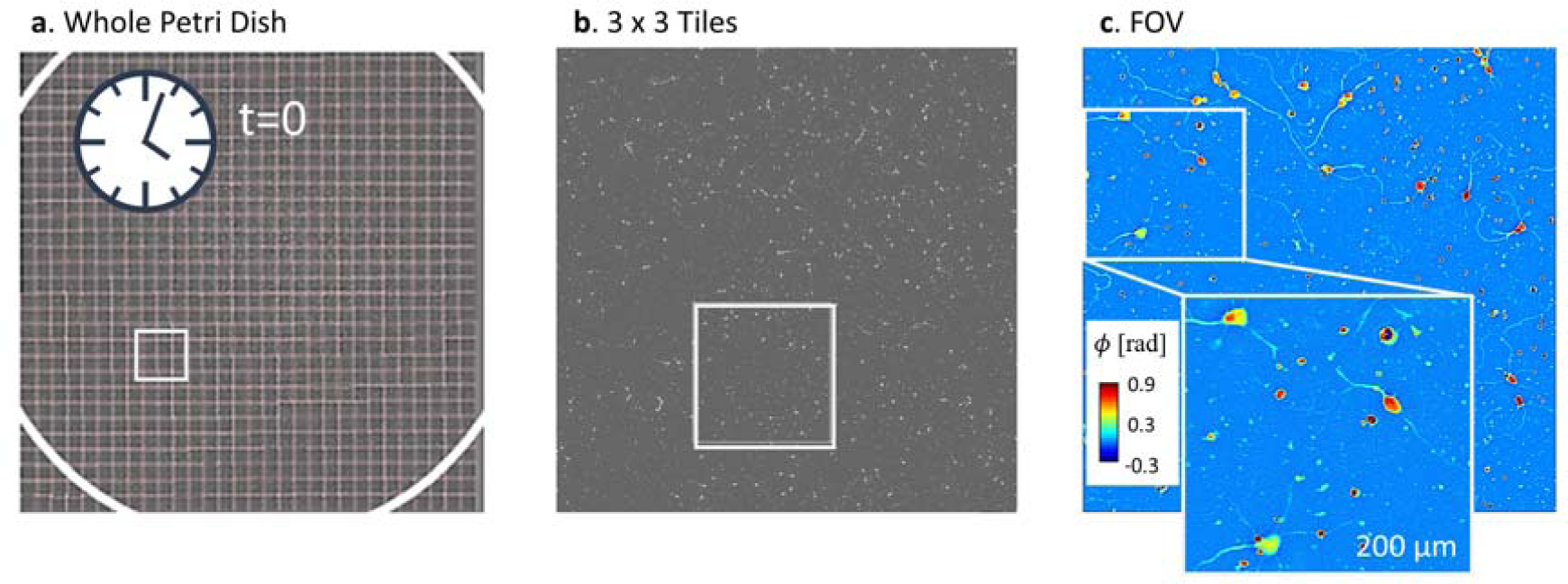
A large field of view assembled from mosaic tiles. (a) To investigate neuron growth we image a whole petri dish (35 mm, 20x/0.3, glass bottom outlined in white) over the course of 29 hours. Each time point is composed of 900 mosaic tiles. Tiles are aligned after assembly with a purpose-built phase correlation algorithm and visualized with TrakEM2. (b,c) zoomed portions of the previous image.

Fig. 7 illustrates the meaning of the artifacts overcome by our technique. As shown in panel A, we correct for the non-physical dip in the phase values near the cell bodies. Indeed, without this correction, much of the mass contribution from short neuronal extensions would be eliminated. Previously, this challenge was mitigated by thresholding out the halo-induced negative values such that no phase shift is allowed to be less than 0 rad[52]. Another category of error occurs when cells cluster together. Similar to the histopathology tissue in Fig. 5, the halo artifact obscures these contact points as well as details within the cell (Fig. 7, “Junctions”), which are important for studying cell-to-cell communication[53].

**Fig. 7.**
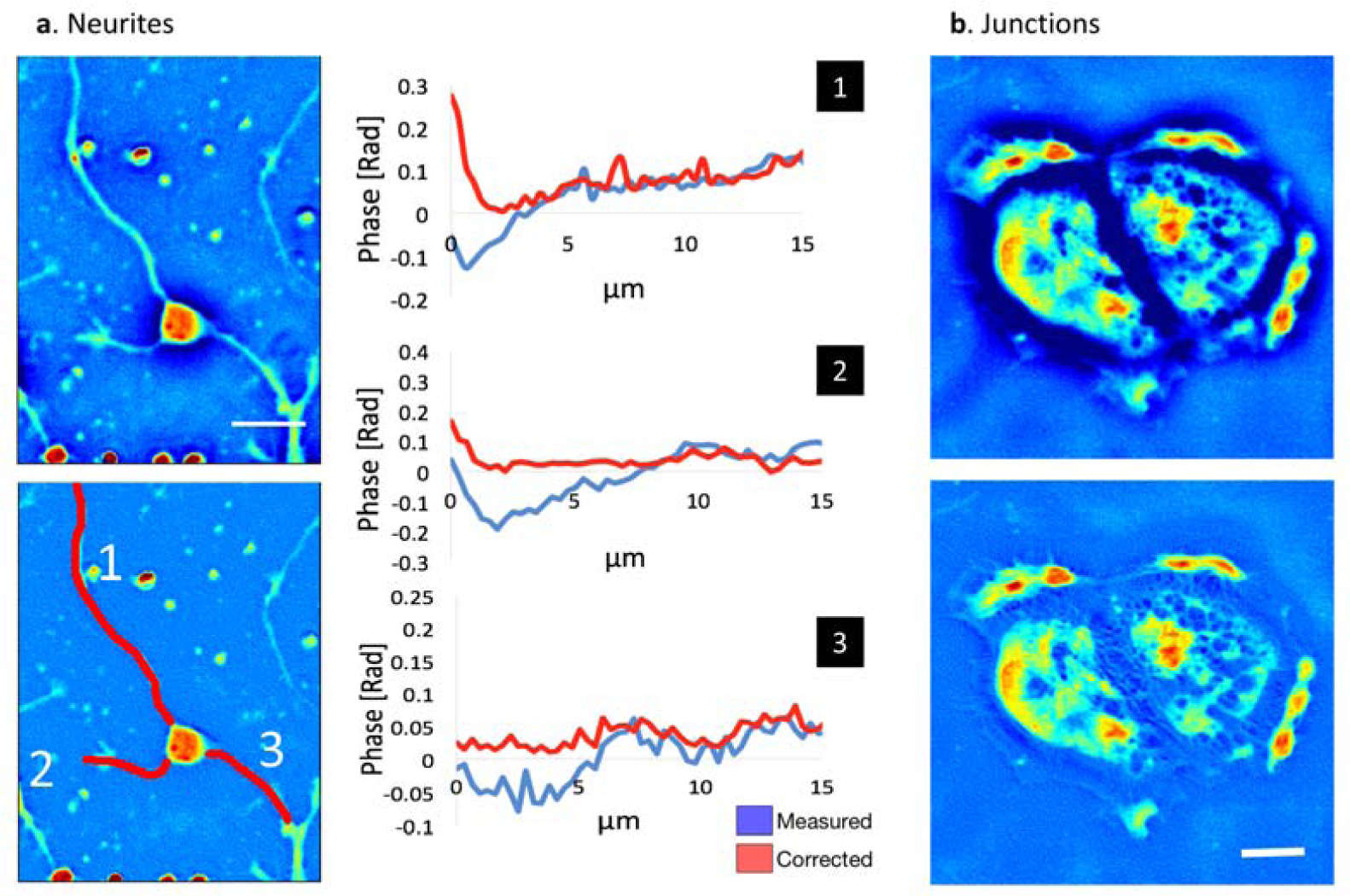
Halo removal fixes neurite near cell bodies and cell clusters. (a) In a typical neuron (white bar = 30 um), the neurites appear submerged under the halo-artifact, showing a non-physical phase shift near the cell body (blue curves). After application of our technique, this defect is removed. (b) Cell clusters (such as the glia shown, white bar = 30 μm) suffer from halo defects, particularly where cells contact each other. After halo removal, similar to tissue in previous figures, a more complex structure becomes visible. Images displayed on a [-0.3,0.9] RAD scale.

As shown in Fig. 8, the halo-free phase maps show an overall positive mass growth rate, while without numerical correction; these images display little to no growth. In contrast, without application of our halo correction routine, it is difficult to detect any mass growth, despite the direct observation of sprouting neurites.

To demonstrate our newfound ability to measure the subtle growth behavior of neuronal cultures, we calculate a histogram of growth rates (Fig. 8(a)). Following the protocol outlined in reference [54], we selected the mosaic tiles within the glass bottom portion of the dish, and obtain a time-lapse growth curve that measures the cellular mass. Under a linear model for cellular growth, the slope of this line gives a growth rate. As expected, after halo removal, the petri dish shows a trend for growth (Fig. 8(b)) as well a small portion of tiles where the overall mass decreased (Fig. 8(c)).

**Fig. 8.**
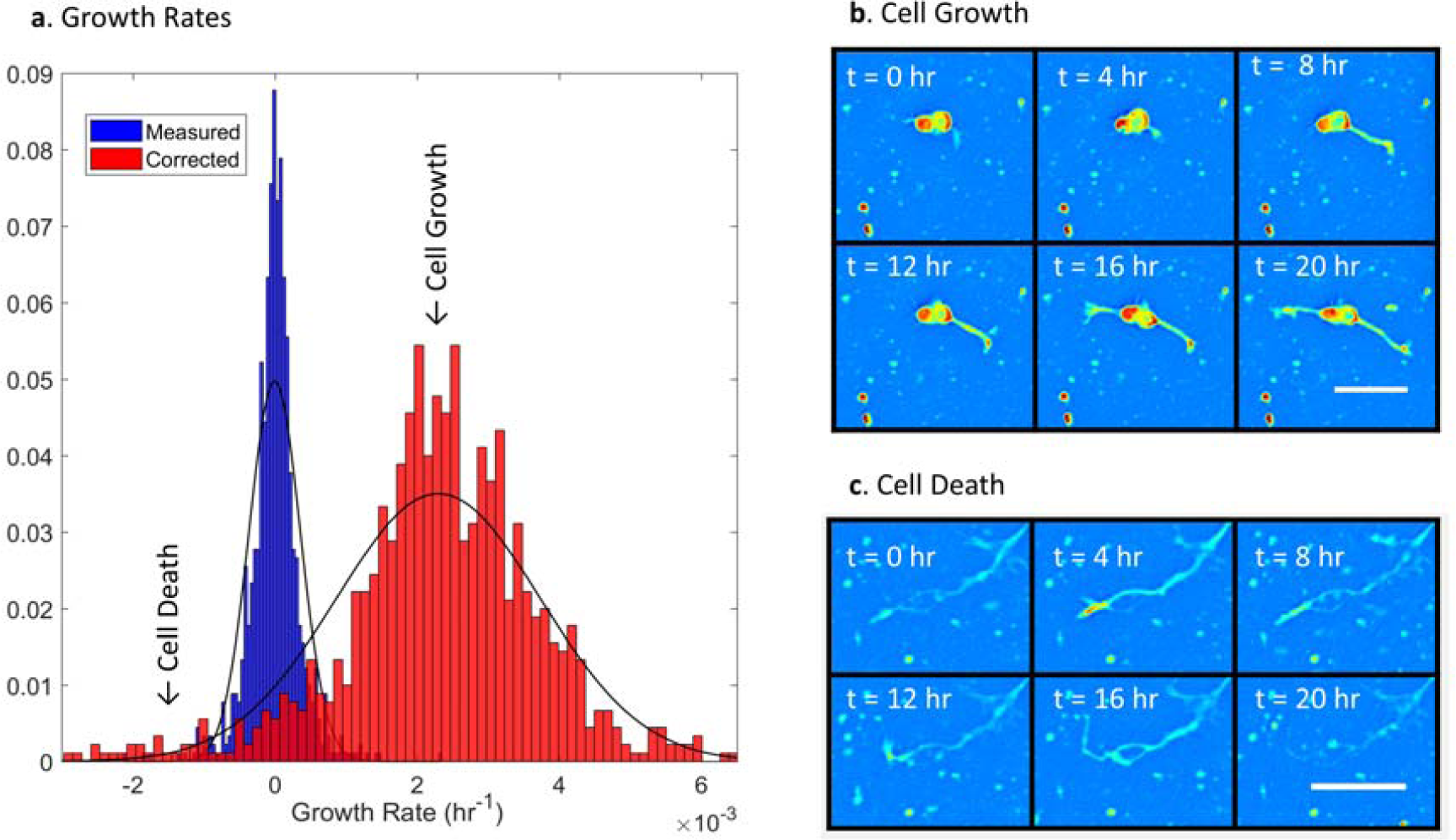
Halo removal reveals cell growth behavior. **(a)** According to a linear growth model, each tile is assigned a rate by finding the change in dry mass over time. The gaussian fit for the growth rate of the measured data has a mean of 0.101 × 10^−3^ hr^−1^ and a standard deviation of 3.6 × 10^−4^ hr^−1^, while that of the corrected data has a mean of 2.41 × 10^−3^ hr^−1^ and a standard deviation of 1.4 ×10^−3^ hr^-1^. Comparing the growth rates between the original and halo-correct data, we observed a mean growth rate shift of 2.3 × 10^−3^ hr^−1^, and calculated a statistically significant difference (p < 0.01). **(b)** In tiles showing positive growth rates, neurite extensions increase in length (representative inset shown), while in the few tiles with negative growth rates, extensions typically wither away (**c**). Images acquired with 20x/0.3 and displayed on a [-0.3,0.9] RAD range with the white bar indicating 20 μm.

Time-lapse sequences of representative fields of view are shown in Visualization 1-3. While the halo-impacted data incorrectly shows that, the cells in the petri dish are not growing. After applying our technique we can accurately study neuronal growth and proliferation.

Neurons exhibit growth rates an order of magnitude smaller than adherent cells, which, due to mitosis, double in mass, roughly, every 10-40 hours (see for points of comparison [55, 56]). In practice, this introduces an instrument sensitivity requirement that is difficult to meet with many methods as imaging artifacts often exceed the 0.15 Rad phase shift of a typical neurite[12, 57]. Comparing to limited previous efforts, the values reported in this work are comparable with those in[58], within variability due to cell type and confluence. One of the few alternatives to optical assays of dry mass involves culturing cells on a purpose-built micro-cantilever device designed to measure mass by detecting changes in resonant frequency. Using such a technique, the authors in [59] reported a growth rate that is, approximately two times larger than what we measured. As those cells are phenotypically closer, we believe that the discrepancy can be attributed to selection bias. Here we measure the growth rates of whole fields of view compared to isolated cells. These fields typically contain over a hundred cells, some of which gain while others lose mass (during cell death, for example). To avoid selection bias, we interferometrically resolved the entire petri dish, to capture the dynamics of the whole population.

## 4. Conclusions

In summary, we present a halo removal technique capable of operating in real-time on existing phase contrast microscopes. Our method relies on using the Hilbert transform to perform integration of the initial phase data along different directions and corroborating those results to preserve the quantitative nature of high-frequency structures. Although the images used in this work were acquired primarily using a SLIM add-on module, we demonstrate that the procedure outlined here can be used in other common-path systems, affected by halos. We show that by correcting the phase values surrounding neuronal extensions, our technique can be used to assay growth and proliferation of cellular cultures in an automated, high-throughput fashion over extended periods without the use of fluorescent markers.

We anticipate that this development will be easy to implement in the microscopy field, as the SLIM module can outfit any phase contrast microscope and the numerical processing runs in real-time. We illustrated the applicability of our method to quantitative studies of sperm cell characterization, red blood cell imaging, tissue diagnosis, and adherent cell growth. However, we expect that halo-free quantitative phase imaging will find numerous other applications, once the technology is adopted broadly by the life science community.

## Acknowledgements

The authors would like to thank Dr. Marcello Rubessa for providing the bovine sperm sample. The tissue microarray was prepared under the supervision of Dr. Andre Kajdacsy-Balla. Aliquots of frozen neurons were obtained from Dr. Jyothi Arikkath at the University of Nebraska Medical Center

## References

1. F. Zernike, “How I Discovered Phase Contrast,” Science 121, 345–349 (1955).

2. E. Abbe, “Beiträge zur Theorie des Mikroskops und der mikroskopischen Wahrnehmung,” (1873).

3. A. F. Huxley and R. Niedergerke, “Structural changes in muscle during contraction; interference microscopy of living muscle fibres,” Nature 173, 971–973 (1954).

4. M. Loehrer, J. Botterweck, J. Jahnke, D. M. Mahlmann, J. Gaetgens, M. Oldiges, R. Horbach, H. Deising, and U. Schaffrath, “In vivo assessment by Mach-Zehnder double-beam interferometry of the invasive force exerted by the Asian soybean rust fungus (Phakopsora pachyrhizi),” New Phytol 203, 620–631 (2014).

5. C. Maurer, A. Jesacher, S. Bernet, and M. Ritsch-Marte, “Phase contrast microscopy with full numerical aperture illumination,” Opt Express 16, 19821–19829 (2008).

6. P. Gao, B. Yao, I. Harder, N. Lindlein, and F. J. Torcal-Milla, “Phase-shifting Zernike phase contrast microscopy for quantitative phase measurement,” Opt Lett 36, 4305–4307 (2011).

7. C. A. Edwards, T. Nguyen, G. Popescu, and L. Goddard, “Image Formation and Halo Removal in Diffraction Phase Microscopy with Partially Coherent Illumination,” in Frontiers in Optics 2014, OSA Technical Digest (online) (Optical Society of America, 2014), FTu4C.5.

8. I. Vartiainen, R. Mokso, M. Stampanoni, and C. David, “Halo suppression in full-field x-ray Zernike phase contrast microscopy,” Opt Lett 39, 1601–1604 (2014).

9. T. Otaki, “Artifact Halo Reduction in Phase Contrast Microscopy Using Apodization,” Optical Review 7, 119–122 (2000).

10. C. Zuo, J. Sun, J. Li, J. Zhang, A. Asundi, and Q. Chen, “High-resolution transport-of-intensity quantitative phase microscopy with annular illumination,” Sci Rep 7, 7654 (2017).

11. G. Popescu, T. Ikeda, R. R. Dasari, and M. S. Feld, “Diffraction phase microscopy for quantifying cell structure and dynamics,” Opt Lett 31, 775–777 (2006).

12. M. Rinehart, Y. Zhu, and A. Wax, “Quantitative phase spectroscopy,” Biomed. Opt. Express 3, 958–965 (2012).

13. H. Farrokhi, J. Boonruangkan, B. J. Chun, T. M. Rohith, A. Mishra, H. T. Toh, H. S. Yoon, and Y. J. Kim, “Speckle reduction in quantitative phase imaging by generating spatially incoherent laser field at electroactive optical diffusers,” Opt Express 25, 10791–10800 (2017).

14. Z. Yin, T. Kanade, and M. Chen, “Understanding the phase contrast optics to restore artifact-free microscopy images for segmentation,” Med Image Anal 16, 1047–1062 (2012).

15. I. Ersoy, F. Bunyak, M. A. Mackey, and K. Palaniappan, “Cell Segmentation Using Hessian-Based Detection and Contour Evolution with Directional Derivatives,” Proc Int Conf Image Proc 2008, 1804–1807 (2008).

16. J. Pan, T. Kanade, and M. Chen, “Learning to Detect Different Types of Cells under Phase Contrast Microscopy,” in Microscopic Image Analysis with Applications in Biology (MIAAB) 2009, 2009),

17. H. Su, Z. Yin, S. Huh, T. Kanade, and J. Zhu, “Interactive Cell Segmentation Based on Active and Semi-Supervised Learning,” IEEE Trans Med Imaging 35, 762–777 (2016).

18. M. E. Ambuhl, C. Brepsant, J. J. Meister, A. B. Verkhovsky, and I. F. Sbalzarini, “High-resolution cell outline segmentation and tracking from phase-contrast microscopy images,” J Microsc 245, 161–170 (2012).

19. N. Jaccard, L. D. Griffin, A. Keser, R. J. Macown, A. Super, F. S. Veraitch, and N. Szita, “Automated method for the rapid and precise estimation of adherent cell culture characteristics from phase contrast microscopy images,” Biotechnol Bioeng 111, 504–517 (2014).

20. T. H. Nguyen, M. Kandel, H. M. Shakir, C. Best-Popescu, J. Arikkath, M. N. Do, and G. Popescu, “Halo-free Phase Contrast Microscopy,” Sci Rep 7, 44034 (2017).

21. G. Popescu, L. P. Deflores, J. C. Vaughan, K. Badizadegan, H. Iwai, R. R. Dasari, and M. S. Feld, “Fourier phase microscopy for investigation of biological structures and dynamics,” Opt Lett 29, 2503–2505 (2004).

22. G. Popescu, Quantitative phase imaging of cells and tissues (McGraw Hill Professional, 2011).

23. P. M. Roma, L. Siman, B. Hissa, U. Agero, E. M. Braga, and O. N. Mesquita, “Profiling of individual human red blood cells under osmotic stress using defocusing microscopy,” J Biomed Opt 21, 90505 (2016).

24. L. Kastl, M. Isbach, D. Dirksen, J. Schnekenburger, and B. Kemper, “Quantitative phase imaging for cell culture quality control,” Cytometry A 91, 470–481 (2017).

25. S. A. Yang, J. Yoon, K. Kim, and Y. Park, “Measurements of morphological and biophysical alterations in individual neuron cells associated with early neurotoxic effects in Parkinson’s disease,” Cytometry A 91, 510–518 (2017).

26. M. E. Kandel, D. Fernandes, A. M. Taylor, H. Shakir, C. Best-Popescu, and G. Popescu, “Three-dimensional intracellular transport in neuron bodies and neurites investigated by label-free dispersion-relation phase spectroscopy,” Cytometry A 91, 519–526 (2017).

27. P. Guo, J. Huang, and M. A. Moses, “Characterization of dormant and active human cancer cells by quantitative phase imaging,” Cytometry A 91, 424–432 (2017).

28. A. Calabuig, M. Mugnano, L. Miccio, S. Grilli, and P. Ferraro, “Investigating fibroblast cells under “safe” and “injurious” blue-light exposure by holographic microscopy,” J Biophotonics 10, 919–927 (2017).

29. T. Lu, B. Corliss, S. Lee, and B. Anvari, “Combined Optical Tweezers and Quantitative Phase Imaging for Mechanical Characterization of Ovarian Cells,” in Optics in the Life Sciences Congress, OSA Technical Digest (online) (Optical Society of America, 2017), OtTu2E.2.

30. A. Fan, A. Tofangchi, M. Kandel, G. Popescu, and T. Saif, “Coupled circumferential and axial tension driven by actin and myosin influences in vivo axon diameter,” Scientific Reports 7, 14188 (2017).

31. A. Badea, J. M. McCracken, E. G. Tillmaand, M. E. Kandel, A. W. Oraham, M. B. Mevis, S. S. Rubakhin, G. Popescu, J. V. Sweedler, and R. G. Nuzzo, “3D-Printed pHEMA Materials for Topographical and Biochemical Modulation of Dorsal Root Ganglion Cell Response,” ACS Applied Materials & Interfaces 9, 30318–30328 (2017).

32. M. Haifler, P. Girshovitz, G. Band, G. Dardikman, I. Madjar, and N. T. Shaked, “Interferometrie phase microscopy for label-free morphological evaluation of sperm cells,” Fertil Steril 104, 43–47 e42 (2015).

33. D. Jin, Y. Sung, N. Lue, Y. H. Kim, P. T. C. So, and Z. Yaqoob, “Large population cell characterization using quantitative phase cytometer,” Cytometry A 91, 450–459 (2017).

34. B. Janicke, A. Karsnas, P. Egelberg, and K. Alm, “Label-free high temporal resolution assessment of cell proliferation using digital holographic microscopy,” Cytometry A 91, 460–469 (2017).

35. P. Cintora, J. Arikkath, M. Kandel, G. Popescu, and C. Best-Popescu, “Cell density modulates intracellular mass transport in neural networks,” Cytometry A 91, 503–509 (2017).

36. H. Majeed, C. Okoro, A. Kajdacsy-Balla, K. C. Toussaint, Jr., and G. Popescu, “Quantifying collagen fiber orientation in breast cancer using quantitative phase imaging,” J Biomed Opt 22, 46004 (2017).

37. T. H. Nguyen, S. Sridharan, V. Macias, A. Kajdacsy-Balla, J. Melamed, M. N. Do, and G. Popescu, “Automatic Gleason grading of prostate cancer using quantitative phase imaging and machine learning,” J Biomed Opt 22, 36015 (2017).

38. M. E. Kandel, K. W. Teng, P. R. Selvin, and G. Popescu, “Label-Free Imaging of Single Microtubule Dynamics Using Spatial Light Interference Microscopy,” ACS Nano 11, 647–655 (2017).

39. H. Schreiber and J. H. Bruning, “Phase Shifting Interferometry,” in Optical Shop Testing (John Wiley & Sons, Inc., 2007), pp. 547–666.

40. B. Bhaduri, H. Pham, M. Mir, and G. Popescu, “Diffraction phase microscopy with white light,” Opt Lett 37, 1094–1096 (2012).

41. T. H. Nguyen, C. Edwards, L. L. Goddard, and G. Popescu, “Quantitative phase imaging with partially coherent illumination,” Opt Lett 39, 5511–5514 (2014).

42. S. Ceballos, M. E. Kandel, S. Sridharan, H. Majeed, F. Monroy, and G. Popescu, “Active intracellular transport in metastatic cells studied by spatial light interference microscopy,” in (SPIE, 2015), 6.

43. R. M. Goldstein, H. A. Zebker, and C. L. Werner, “Satellite radar interferometry: Two-dimensional phase unwrapping,” Radio Science 23, 713–720 (1988).

44. M. E. Kandel, S. Sridharan, J. Liang, Z. Luo, K. Han, V. Macias, A. Shah, R. Patel, K. Tangella, A. Kajdacsy-Balla, G. Guzman, and G. Popescu, “Label-free tissue scanner for colorectal cancer screening,” J Biomed Opt 22, 66016 (2017).

45. H. V. Pham, B. Bhaduri, K. Tangella, C. Best-Popescu, and G. Popescu, “Real time blood testing using quantitative phase imaging,” PLoS One 8, e55676 (2013).

46. M. Takeda, H. Ina, and S. Kobayashi, “Fourier-transform method of fringe-pattern analysis for computer-based topography and interferometry,” Journal of the Optical Society of America 72, 156–160 (1982).

47. T. Ikeda, G. Popescu, R. R. Dasari, and M. S. Feld, “Hilbert phase microscopy for investigating fast dynamics in transparent systems,” Opt Lett 30, 1165–1167 (2005).

48. S. Wang, K. Yan, and L. Xue, “Quantitative interferometric microscopy with two dimensional Hilbert transform based phase retrieval method,” Optics Communications 383, 537–544 (2017).

49. M. R. Arnison, C. J. Cogswell, N. I. Smith, P. W. Fekete, and K. G. Larkin, “Using the Hilbert transform for 3D visualization of differential interference contrast microscope images,” Journal of Microscopy 199, 79–84 (2000).

50. M. R. Arnison, K. G. Larkin, C. J. Sheppard, N. I. Smith, and C. J. Cogswell, “Linear phase imaging using differential interference contrast microscopy,” J Microsc 214, 7–12 (2004).

51. C. A, “TrakEM2: an ImageJ-based program for morphological data mining and 3D modeling,” in Proceedings of the 1st ImageJ User and Developer Conference, 2006), 18–19.

52. M. Mir, Z. Wang, Z. Shen, M. Bednarz, R. Bashir, I. Golding, S. G. Prasanth, and G. Popescu, “Optical measurement of cycle-dependent cell growth,” Proc Natl Acad Sei U S A 108, 13124–13129 (2011).

53. H. Delanoe-Ayari, P. Lenz, J. Brevier, M. Weidenhaupt, M. Vallade, D. Gulino, J. F. Joanny, and D. Riveline, “Periodic adhesive fingers between contacting cells,” Phys Rev Lett 93, 108102 (2004).

54. G. Popescu, Y. Park, N. Lue, C. Best-Popescu, L. Deflores, R. R. Dasari, M. S. Feld, and K. Badizadegan, “Optical imaging of cell mass and growth dynamics,” Am J Physiol Cell Physiol 295, C538–544 (2008).

55. M. Mir, A. Bergamaschi, B. S. Katzenellenbogen, and G. Popescu, “Highly sensitive quantitative imaging for monitoring single cancer cell growth kinetics and drug response,” PLoS One 9, e89000 (2014).

56. T. H. Nguyen, M. E. Kandel, M. Rubessa, M. B. Wheeler, and G. Popescu, “Gradient light interference microscopy for 3D imaging of unlabeled specimens,” Nat Commun 8, 210 (2017).

57. R. Schubert, A. Vollmer, S. Ketelhut, and B. Kemper, “Enhanced quantitative phase imaging in self-interference digital holographic microscopy using an electrically focus tunable lens,” Biomed. Opt. Express 5, 4213–4222 (2014).

58. M. Mir, T. Kim, A. Majumder, M. Xiang, R. Wang, S. C. Liu, M. U. Gillette, S. Stice, and G. Popescu, “Label-free characterization of emerging human neuronal networks,” Sci Rep 4, 4434 (2014).

59. E. A. Corbin, L. J. Millet, K. R. Keller, W. P. King, and R. Bashir, “Measuring physical properties of neuronal and glial cells with resonant microsensors,” Anal Chem 86, 4864–4872 (2014).

